# Tree species natural regeneration in a tidal white-water floodplain forest of the Amazon River estuary

**DOI:** 10.1101/2025.10.03.680168

**Authors:** Caroline da Cruz Vasconcelos, Glaucileide Ferreira, Camila Brandão da Silva, Cryslene Furtado da Costa, Jaynna Gonar Lôbo Isacksson, Marciane Furtado Freitas, Noelle Loyanna Lima Almeida, Vanessa Carla Campelo de Sousa, Laura Lorena Rivera-Parada, Kaio Cesar Marinho da Cunha, Semírian Campos Amoêdo, Gabrielly Guabiraba-Ribeiro, Adelson Rocha Dantas, Robson Borges de Lima, Jadson Coelho de Abreu

## Abstract

The Amazon hosts the world’s most diverse floodplain forests, characterized by tree communities adapted to seasonal flooding. In the estuary, these forests are driven by both fluvial and tidal dynamics, yet remain poorly studied. We assessed natural regeneration in a tidal white-water floodplain forest within the Fazendinha Environmental Protection Area, eastern Amazonia (Brazil). Regenerating individuals (DSH ≤ 5 cm) were sampled in 21 systematically distributed plots, classified into height classes, and analyzed with diversity indices, regeneration parameters, and ordination methods. We recorded 758 individuals from 43 taxa, 35 genera, and 20 families. Fabaceae was the richest and most abundant family, largely due to the oligarch *Mora paraensis*, which comprised >70% of individuals. Despite this dominance, Shannon diversity was high (H’ = 5.31), with 53.5% of species occurring as singletons or doubletons. Regeneration density reached 14,438 ind. ha□^1^, concentrated in the smallest classes, with few species spanning all classes. Subtle compositional shifts occurred with river distance, though dominants remained broadly distributed. Our findings highlight the resilience of oligarchic species and the vulnerability of rare taxa, advancing knowledge of estuarine floodplain ecology and providing essential baseline information for conservation and management in this understudied region.

## Introduction

Tropical forests play a fundamental role in maintaining global biodiversity and regulating ecosystem services (Cao and Woodward 1998; Artaxo et al. 2022). Among them, the Amazon hosts the highest tree diversity on Earth, harboring more than one-third of all Neotropical species (Gentry 1982; Antonelli et al. 2011). This diversity is distributed across a mosaic of habitats shaped by heterogeneous soils, topography, and hydrological regimes (Pitman et al. 2001; Fine et al. 2005; Quesada et al. 2012; Schietti et al. 2014; Tuomisto et al. 2017). Amazonian floodplains are home to complex ecosystems, where periodic inundation acts as a strong ecological filter, favoring species capable of tolerating prolonged waterlogging and sediment deposition (Wittmann et al. 2004; Parolin 2009). These habitats are therefore hotspots of ecological specialization (Wittmann et al. 2013), yet they remain underrepresented in floristic and ecological surveys, particularly in the estuarine portion of the Amazon Basin.

Floodplain forests associated with white-water rivers (also called *várzea* forest) are among the most productive ecosystems in the tropics (Haugaasen and Peres 2006; Petsch et al. 2023). They cover ∼516,000 km^2^ of the Amazon (Hess et al. 2015) and support approximately one-sixth of its tree flora (Householder et al. 2024). Their high fertility, dynamic hydrology, and geomorphological processes generate complex patterns of species dominance, evenness, and turnover, resulting in multiple forest communities that differ in age, physiognomy, and floristic composition (Wittmann and Junk 2003). In many cases, a few dominant taxa often account for the majority of individuals, while many others persist as rare species (Whittaker 1965; ter Steege et al. 2013). Understanding these patterns is crucial, as species dominance not only shapes community structure and ecosystem functioning but also influences ecosystem resilience to environmental change and human disturbance. However, most available knowledge derives from central Amazonian white-water floodplain forests (Wittmann and Junk 2003; Oliveira-Wittmann et al. 2007; Assis and Wittmann 2011), leaving estuarine floodplains comparatively understudied.

The Amazon estuary represents an ecotonal region where riverine flooding interacts with tidal oscillations, producing two daily flood pulses and steep environmental gradients over short spatial scales (Junk et al. 2011). These conditions strongly influence seed dispersal, establishment, and regeneration, making estuarine white-water floodplain forests ideal systems to investigate how hydrological regimes filter species composition. Despite their ecological and socio-economic importance, however, estuarine white-water floodplain forests remain poorly documented (Carvalho et al. 2023), and baseline data on tree regeneration are virtually absent for the eastern Amazonia, especially in peri-urban landscapes where conservation challenges are pronounced.

Tree regeneration is a key demographic process linking current community composition to future forest structure (Swaine and Whitmore 1988; Díaz-Yáñez et al. 2024). Here, we adopt the concept of “natural regeneration” as summarized by Jardim (2015), which defines it as the quantification of a stand or population at a given moment based on pre-established criteria. Natural regeneration results from interactions among ecological processes that restore forest ecosystems, representing the initial phases of establishment and development within the forest growth cycle (Gama et al. 2002).

Assessing the diversity, abundance, and distribution of juvenile individuals across regeneration classes provides critical insights into species’ potential for persistence and recruitment into the canopy (Assis and Wittmann 2011). In floodplain systems, where hydrological constraints can create demographic bottlenecks, regeneration studies are especially valuable for identifying species resilient to disturbance as well as those vulnerable to local decline (Rother et al. 2013). Moreover, regeneration data can reveal whether canopy dominants maintain strong recruitment (reinforcing oligarchic patterns) or whether rare taxa contribute disproportionately to forest renewal.

In this study, we provide the first community-level assessment of natural regeneration in a white-water floodplain forest influenced by tidal dynamics in the Amazon River estuary. Specifically, we aimed to: (i) describe the floristic composition and diversity of regeneration; (ii) assess regeneration structure across height classes; and (iii) test whether species composition varies along an environmental gradient represented by distance from the river. By addressing this knowledge gap in the eastern Amazonia, particularly in the state of Amapá, our findings provide critical insights for biodiversity conservation, forest management, and the evaluation of estuarine floodplain resilience.

## Material and methods

### Study area

We carried out the study in the Fazendinha Environmental Protection Area (51°07′41.78′′ W and 00°03′10.29′′ S; hereafter APA Fazendinha), a conservation unit that covers an area of 136.59 ha. The APA Fazendinha is located in the peri-urban area between the municipalities of Macapá and Santana, in the state of Amapá, eastern Brazilian Amazonia (Fig. 1). The white-water river floodplain under study is influenced by the tidal dynamics of the Amazon River, characterized by the occurrence of two daily flood pulses (Junk et al. 2011). Flood water can reach heights of up to 60 cm on tree trunks within the forest (Dantas et al. 2021) or up to 4 m during the rainy season at the Amazon River delta (Cunha et al. 2017).

**Fig. 1.**
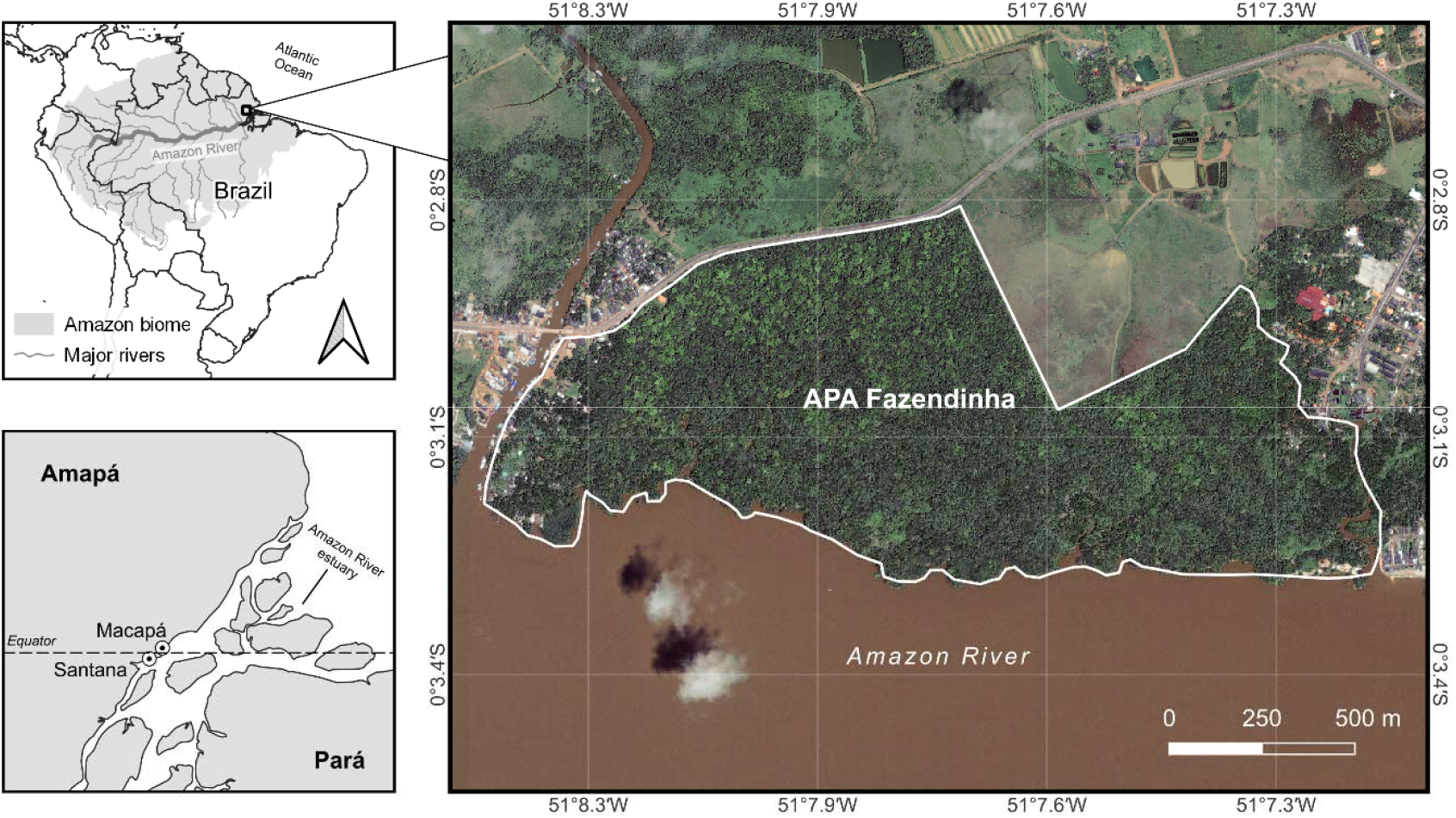
Location of study area in the state of Amapá, eastern Brazilian Amazonia. Map created with QGIS software (v.3.28.1; QGIS Development Team 2022).

The regional climate corresponds to the Am type (rainy tropical) according to Köppen’s classification (Alvares et al. 2013), with the following average annual values: precipitation of 2,549.7 mm, temperature between 23.8 and 31.5 °C, and relative humidity around 82.2% (INMET 2018). The highest rainfall occurs between January and June, and relatively less rainfall is observed in the other months of the year (INMET 2018). The predominant soil is Typical Eutrophic Ta Melanic Gleysol, shallow, silty, and fertile, which may show some acidity, toxicity, and deficiency of certain nutrients (Pinto 2014). The primary vegetation type in APA Fazendinha is white-water flooded forest, formally classified as Alluvial Dense Ombrophilous Forest by IBGE (2012), with elevations ranging between 5 and 26 m.

### Sampling procedures

Individuals of tree regeneration (seedlings and saplings) with a diameter at soil height (DSH) ≤ 5 cm were recorded in twenty-one plots (sample units of 5 × 5 m), systematically distributed along three linear transects perpendicular to the mainstem of the Amazon River. The transects and plots were separated by at least 400 m and 40 m from one another, respectively. The center of each plot was georeferenced using a Garmin^®^ 76CSx GPS. Individuals of tree regeneration were divided into four classes according to height (adapted from Volpato (1994): class 1 (0.5 ≤ height < 1.0 m), class 2 (1.0 ≤ height < 2.0 m), class 3 (2.0 ≤ height < 3 m), and class 4 (height ≥ 3.0 m). DSH and height (h) were field-measured using a digital caliper (Carbografite^®^ model 150) and a tape measure attached to a wooden stick, respectively.

Taxonomic identification was carefully conducted in the field with assistance from a trained parabotanist (a local person knowledgeable on the regional flora) and/or using specialized field guides (e.g., Ribeiro et al. 1999; Camargo et al. 2008; Wittmann et al. 2010). Individuals not identified at the species level were assigned as morphospecies to deal with taxonomic constraints.

### Data preparation and analysis

Scientific names were standardized using the web-based tool Taxonomic Name Resolution Service (TNRS; https://tnrs.biendata.org/), with parameters set to a minimum match threshold of 0.85, World Flora Online (WFO) for family classification, and global taxonomic sources. Order-level classifications followed the Angiosperm Phylogeny Group IV (APG IV 2016).

To characterize the diversity of the regenerating tree assemblage, we applied interpolation (rarefaction) and extrapolation (estimation) curves based on the first three Hill numbers: q = 0 (species richness), q = 1 (the exponential of Shannon’s entropy index), and q = 2 (the inverse of Simpson’s concentration index) (Chao et al. 2014). Confidence intervals (95%) were constructed using 500 bootstraps. The Chao method and Hill numbers are widely used to reduce the effect of insufficient sampling effort in the quantification of diversity (Gotelli et al. 2012; Chao et al. 2014). In addition, we used the rank-abundance plot (Whittaker 1965) to visualize species abundance distribution, as it enhances clarity when displaying relative abundances of a few species compared to histograms (Wilson 1991; Magurran 2004).

We calculated the natural regeneration parameters based on absolute and relative values of Density (D) and Frequency (F), which were also used to calculate the Natural Regeneration per Class (NRC) and the Total Natural Regeneration (TNR) adapted from Volpato (1994): *NRc*_*ij*_ = (*rD*_*ij*_ *+ rF*_*ij*_)⁄2 and *TNR*_*i*_ = ∑(*NRc*_*ij*_)⁄4, where NRC_ij_, natural regeneration of ith species in the jth height class; rD_ij_, the relative density of ith species in the jth height class; rF_ij_, the relative frequency of ith species in the jth height class; and TNR_i_, total natural regeneration of sampled population of ith species. For all families, we determined Family Importance Value (FIV), which is the sum of relative density, relative dominance, and relative diversity, where relative diversity is the number of species of family i/total number of species (Mori et al. 1983).

We built a matrix of species (columns) per plot (rows) for each quantitative (relative abundance) and qualitative (presence-absence) dataset. We included both data types to understand the contribution of abundant and rare species to the distribution pattern (Zuquim et al. 2009). Principal Coordinate Analysis (PCoA) with Cailliez correction was performed to reduce the dimensionality of the species composition (assemblage structure). In the first ordination, we applied the Bray-Curtis dissimilarity index using the Hellinger-transformed species abundance data. In the second ordination, we applied the Sørensen dissimilarity index using species presence-absence data. In both cases, we used the first axis derived from PCoA to test the relationship between the response variable (PCoA-1) and the environmental gradient by performing a simple regression. The distance from the Amazon River mainstem was used as a proxy to represent the environmental gradient along plots. We measured the distance (in meters) from the center of each sample plot to the Amazon River.

Unless indicated, all analyses and figures were performed using the open-source statistical software R (v.4.4.2; R Core Team 2024), employing a custom-made R script and the add-on packages listed in Supplementary Information Table S1.

## Results

### Floristic diversity of tree species natural regeneration

We recorded a total of 758 individuals (DSH ≤ 5 cm) across the study area, comprising 20 botanical families, 35 genera, and 43 taxa (diagnosable species and morphotypes) (Table 1). Among these taxa, 13 were identified at the genus level and one at the family level. In terms of number of species, the richest families were Fabaceae (12 spp.), followed by Bignoniaceae, Sapotaceae, and Violaceae in the same rank (3 spp. each). In order level, the best-represented taxa in the community were Fabales and Malpighiales, and the phylogenetic relationships of the local species pool can be consulted in Supplementary Information Fig. S2.

**Table 1.**
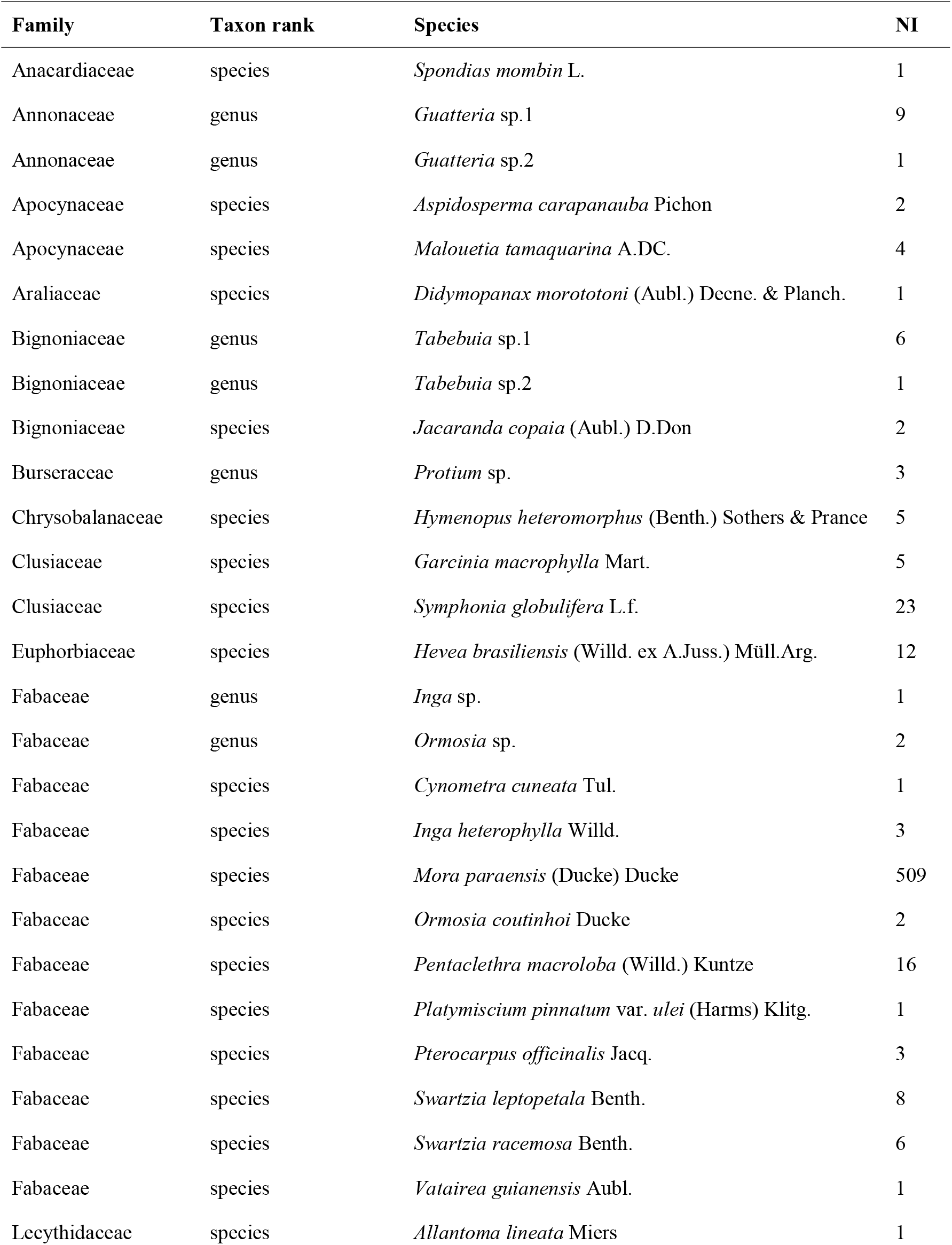
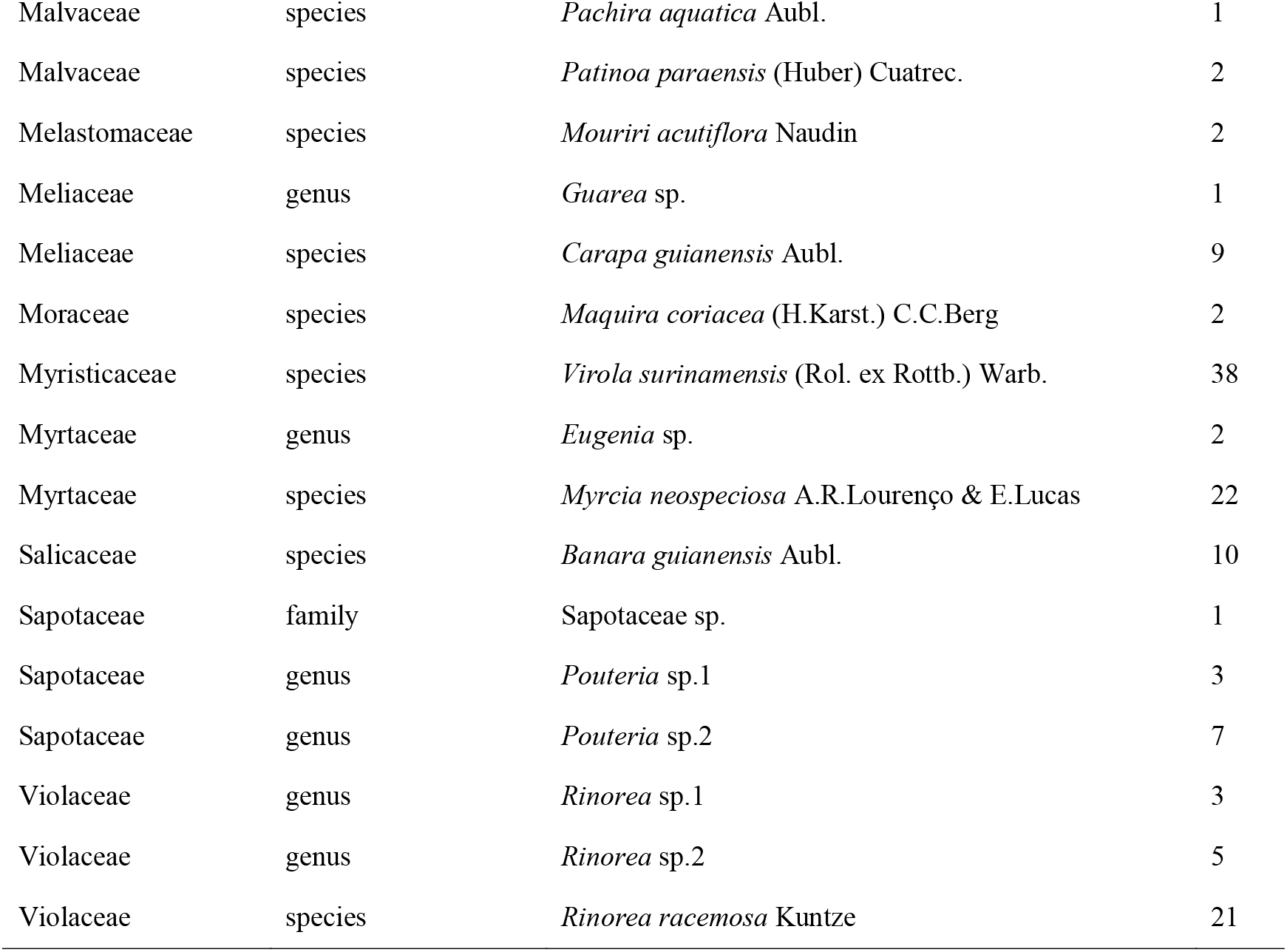
Floristic composition of the tree species natural regeneration in a tidal white-water floodplain forest of the Amazon River, eastern Amazonia. List of families (in alphabetical order), followed by taxon rank, scientific name, and their respective number of individuals (NI).

| Family | Taxon rank | Species | NI |
| --- | --- | --- | --- |
| Anacardiaceae | species | *Spondias mombin* L. | 1 |
| Annonaceae | genus | *Guatteria* sp.1 | 9 |
| Annonaceae | genus | *Guatteria* sp.2 | 1 |
| Apocynaceae | species | *Aspidosperma carapanauba* Pichon | 2 |
| Apocynaceae | species | *Malouetia tamaquarina* A.DC. | 4 |
| Araliaceae | species | *Didymopanax morototoni* (Aubl.) Decne. & Planch. | 1 |
| Bignoniaceae | genus | *Tabebuia* sp.1 | 6 |
| Bignoniaceae | genus | *Tabebuia* sp.2 | 1 |
| Bignoniaceae | species | *Jacaranda copaia* (Aubl.) D.Don | 2 |
| Burseraceae | genus | *Protium* sp. | 3 |
| Chrysobalanaceae | species | *Hymenopus heteromorphus* (Benth.) Sothers & Prance | 5 |
| Clusiaceae | species | *Garcinia macrophylla* Mart. | 5 |
| Clusiaceae | species | *Symphonia globulifera* L.f. | 23 |
| Euphorbiaceae | species | *Hevea brasiliensis* (Willd. ex A.Juss.) Müll.Arg. | 12 |
| Fabaceae | genus | *Inga* sp. | 1 |
| Fabaceae | genus | *Ormosia* sp. | 2 |
| Fabaceae | species | *Cynometra cuneata* Tul. | 1 |
| Fabaceae | species | *Inga heterophylla* Willd. | 3 |
| Fabaceae | species | *Mora paraensis* (Ducke) Ducke | 509 |
| Fabaceae | species | *Ormosia coutinhoi* Ducke | 2 |
| Fabaceae | species | *Pentaclethra macroloba* (Willd.) Kuntze | 16 |
| Fabaceae | species | *Platymiscium pinnatum* var. *ulei* (Harms) Klitg. | 1 |
| Fabaceae | species | *Pterocarpus officinalis* Jacq. | 3 |
| Fabaceae | species | *Swartzia leptopetala* Benth. | 8 |
| Fabaceae | species | *Swartzia racemosa* Benth. | 6 |
| Fabaceae | species | *Vatairea guianensis* Aubl. | 1 |
| Lecythidaceae | species | *Allantoma lineata* Miers | 1 |
| Malvaceae | species | *Pachira aquatica* Aubl. | 1 |
| Malvaceae | species | *Patinoa paraensis* (Huber) Cuatrec. | 2 |
| Melastomaceae | species | *Mouriri acutiflora* Naudin | 2 |
| Meliaceae | genus | *Guarea* sp. | 1 |
| Meliaceae | species | *Carapa guianensis* Aubl. | 9 |
| Moraceae | species | *Maquira coriacea* (H.Karst.) C.C.Berg | 2 |
| Myristicaceae | species | *Virola surinamensis* (Rol. ex Rottb.) Warb. | 38 |
| Myrtaceae | genus | *Eugenia* sp. | 2 |
| Myrtaceae | species | *Myrcia neospeciosa* A.R.Lourenço & E.Lucas | 22 |
| Salicaceae | species | *Banara guianensis* Aubl. | 10 |
| Sapotaceae | family | Sapotaceae sp. | 1 |
| Sapotaceae | genus | *Pouteria* sp.1 | 3 |
| Sapotaceae | genus | *Pouteria* sp.2 | 7 |
| Violaceae | genus | *Rinorea* sp.1 | 3 |
| Violaceae | genus | *Rinorea* sp.2 | 5 |
| Violaceae | species | *Rinorea racemosa* Kuntze | 21 |

Regarding species diversity (Fig. 2), the rarefaction/extrapolation curve based on the richness estimator suggests that, with a more intensive sampling effort, we could identify at least nine additional species. Furthermore, it did not estimate a tendency for the curve to stabilize at an asymptote. On the other hand, the Shannon and Simpson estimators indicate that the diversity recorded adequately reflects the community studied. This is evidenced by the rapid stabilization of the rarefaction/extrapolation curves and the proximity between the observed and estimated values.

**Fig. 2.**
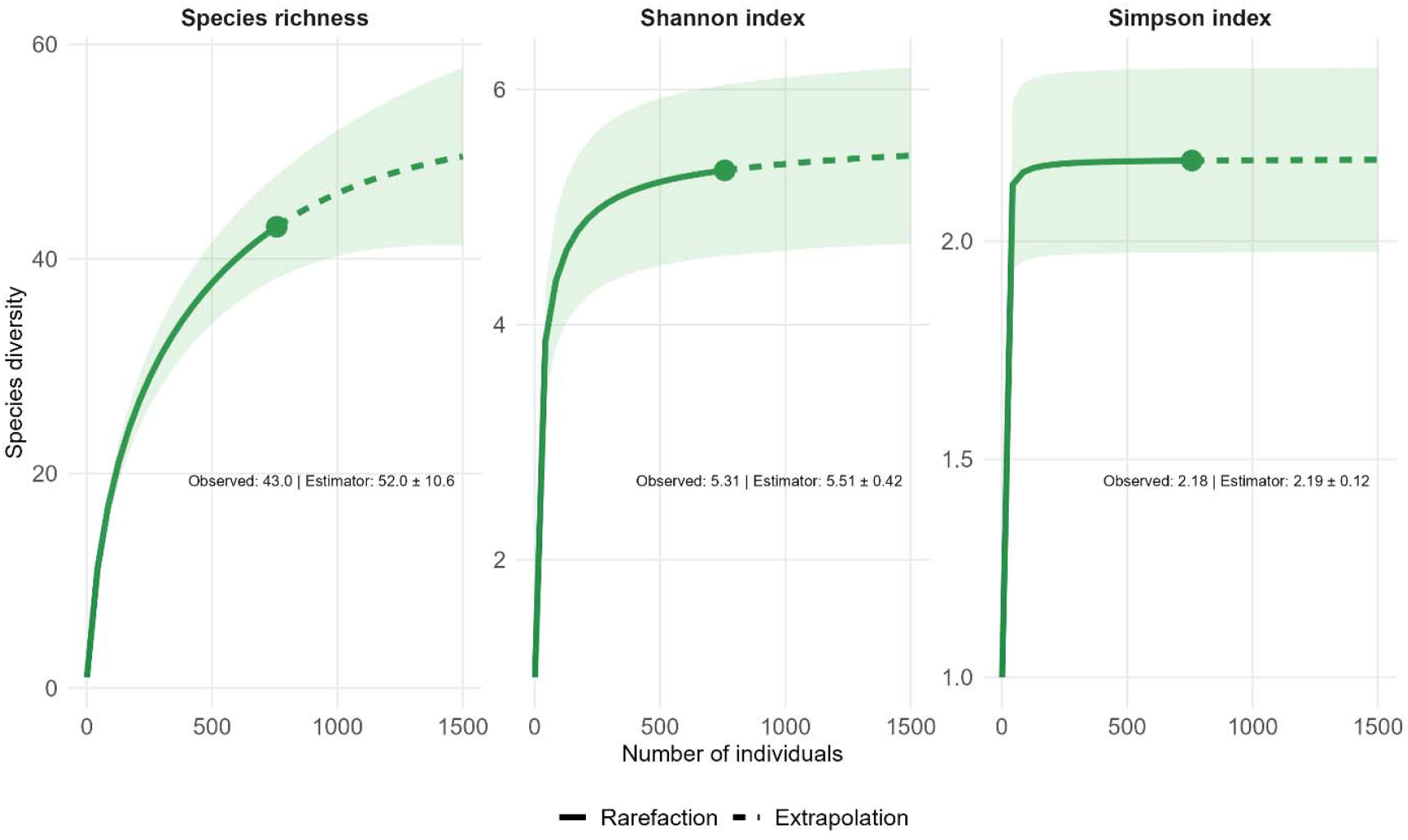
Rarefaction (solid line) and extrapolation (dashed line) sampling curves for three Hill numbers, with corresponding 95% confidence intervals (shaded area; 500 bootstraps), based on individual-based abundance data. The observed diversity (solid circles) and asymptotic estimates with their standard errors are shown in each graph for the regenerating assemblage of tree species in a tidal white-water floodplain forest of the Amazon River estuary, eastern Amazonia.

The five most important families accounted for 46.5% of all sampled species and 88.6% of all sampled individuals. Fabaceae was the most important family (IVF 165.2%), followed by Violaceae (11%), Myrtaceae (14.8%), Myristicaceae (11%), and Clusiaceae (10.2%) (Fig. 3a). Fabaceae had the highest number of individuals (73% of total), followed by Myristicaceae (5%), Violaceae (3.8%), Clusiaceae (3.7%), and Myrtaceae (3.2%) families.

**Fig. 3.**
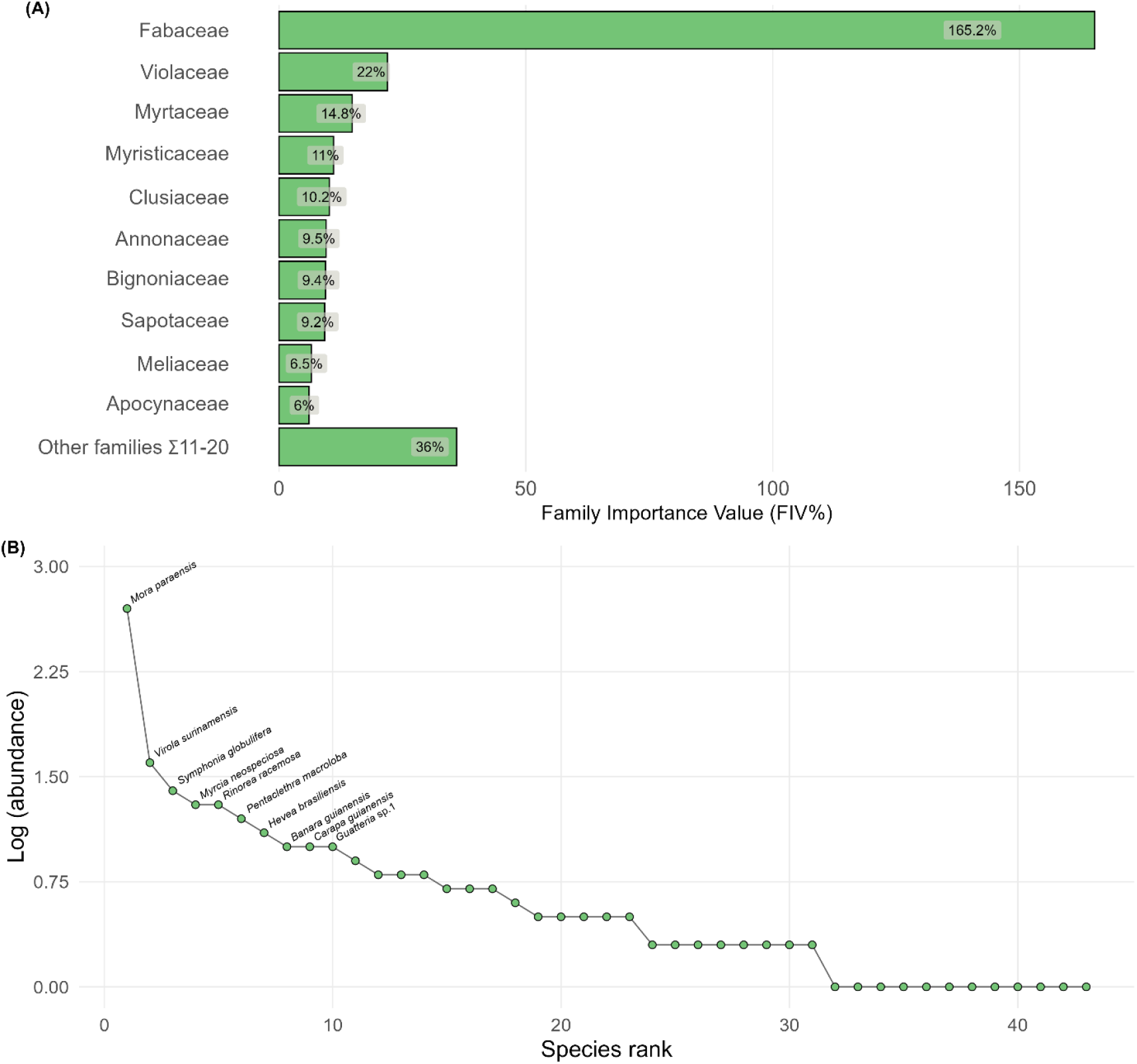
Ten most important families (a) in decreasing order of importance value, including the values summed from remaining families. Rank-abundance plot (b) based on absolute abundance on a logarithmic scale, highlighting the five most abundant species in the regenerating assemblage of tree species in a tidal white-water floodplain forest of the Amazon River estuary, eastern Amazonia.

The overall most abundant species (Fig. 3b) were *Mora paraensis* (509 individuals), which also contributed to the high representativeness of the Fabaceae family (Fig. 2a). Subsequently are *Virola surinamensis* (38 ind.), *Symphonia globulifera* (23 ind.), *Myrcia neospeciosa* (22 ind.), *Rinorea racemosa* (21 ind.), and *Pentaclethra macroloba* (16 ind.), which together account for 83% of sampled individuals. *Mora paraensis* was recorded in the highest number of sample plots (17 plots, 81% of the total sampled), followed by *Symphonia globulifera* and *Myrcia neospeciosa* (12 plots each, 57.1%), as well as *Rinorea racemosa* (10 plots, 47.6%), *Virola surinamensis* (9 plots, 42.9%), and *Pentaclethra macroloba* (8 plots, 38%). In contrast, a large proportion of species (23 species, 53.5% of the total sampled) occurred in at least two sample plots. Singletons and doubletons (*sensu* Magurran 2004) were represented by 14 and 9 species, respectively, together accounting for 53.5% of total species richness.

### Structure of tree species natural regeneration

The previous results are further supported by the Total Natural Regeneration (TNR) rates, which showed *Mora paraensis* as the most representative tree species of the regenerating assemblage. An overview of ten species with the highest TNR is shown in Fig. 4, and a complete list is provided in Table 2. *Mora paraensis, Rinorea racemosa*, and *Myrcia neospeciosa* represented the highest rates of TNR in APA Fazendinha.

**Table 2.** Regenerating assemblage of tree species in a tidal white-water floodplain forest of the Amazon River estuary, eastern Amazonia, listed in decreasing order of TNR. Abbreviations: rF (relative Frequency), rD (relative Density), NRC (Natural Regeneration per Class), and TNR (Total Natural Regeneration).

| Species | Class 1 |  |  | Class 2 |  |  | Class 3 |  |  | Class 4 |  |  | TNR |
| --- | --- | --- | --- | --- | --- | --- | --- | --- | --- | --- | --- | --- | --- |
|  | rF1 | rD1 | NRC1 | rF2 | rD2 | NRC2 | rF3 | rD3 | NRC3 | rF4 | rD4 | NRC4 |  |
| *Mora paraensis* | 12.4 | 66.6 | 39.5 | 26.42 | 74.18 | 50.3 | 35 | 56.67 | 45.83 | 0 | 0 | 0 | 33.91 |
| *Rinorea racemosa* | 4.96 | 1.68 | 3.32 | 7.55 | 2.75 | 5.15 | 25 | 16.67 | 20.83 | 22.22 | 20 | 21.11 | 12.6 |
| *Myrcia neospeciosa* | 6.61 | 2.8 | 4.71 | 3.77 | 1.65 | 2.71 | 5 | 3.33 | 4.17 | 33.33 | 30 | 31.67 | 10.81 |
| *Guatteria* sp.1 | 0.83 | 0.75 | 0.79 | 3.77 | 1.1 | 2.44 | 10 | 6.67 | 8.33 | 11.11 | 10 | 10.56 | 5.53 |
| *Swartzia racemosa* | 2.48 | 0.56 | 1.52 | 0 | 0 | 0 | 5 | 3.33 | 4.17 | 11.11 | 20 | 15.56 | 5.31 |
| *Rinorea* sp.2 | 2.48 | 0.56 | 1.52 | 0 | 0 | 0 | 5 | 3.33 | 4.17 | 11.11 | 10 | 10.56 | 4.06 |
| *Virola surinamensis* | 6.61 | 5.41 | 6.01 | 9.43 | 4.95 | 7.19 | 0 | 0 | 0 | 0 | 0 | 0 | 3.3 |
| *Didymopanax morototoni* | 0 | 0 | 0 | 0 | 0 | 0 | 0 | 0 | 0 | 11.11 | 10 | 10.56 | 2.64 |
| *Symphonia globulifera* | 9.92 | 3.92 | 6.92 | 1.89 | 1.1 | 1.49 | 0 | 0 | 0 | 0 | 0 | 0 | 2.1 |
| *Pentaclethra macroloba* | 4.96 | 2.43 | 3.69 | 5.66 | 1.65 | 3.65 | 0 | 0 | 0 | 0 | 0 | 0 | 1.84 |
| *Hevea brasiliensis* | 4.13 | 2.05 | 3.09 | 0 | 0 | 0 | 5 | 3.33 | 4.17 | 0 | 0 | 0 | 1.81 |
| *Tabebuia* sp.1 | 2.48 | 0.75 | 1.61 | 1.89 | 0.55 | 1.22 | 5 | 3.33 | 4.17 | 0 | 0 | 0 | 1.75 |
| *Swartzia leptopetala* | 3.31 | 0.75 | 2.03 | 7.55 | 2.2 | 4.87 | 0 | 0 | 0 | 0 | 0 | 0 | 1.72 |
| *Inga heterophylla* | 0.83 | 0.19 | 0.51 | 1.89 | 0.55 | 1.22 | 5 | 3.33 | 4.17 | 0 | 0 | 0 | 1.47 |
| *Banara guianensis* | 3.31 | 1.49 | 2.4 | 3.77 | 1.1 | 2.44 | 0 | 0 | 0 | 0 | 0 | 0 | 1.21 |
| *Carapa guianensis* | 4.13 | 1.49 | 2.81 | 1.89 | 0.55 | 1.22 | 0 | 0 | 0 | 0 | 0 | 0 | 1.01 |
| *Garcinia macrophylla* | 2.48 | 0.56 | 1.52 | 3.77 | 1.1 | 2.44 | 0 | 0 | 0 | 0 | 0 | 0 | 0.99 |
| *Pouteria* sp.2 | 1.65 | 0.93 | 1.29 | 3.77 | 1.1 | 2.44 | 0 | 0 | 0 | 0 | 0 | 0 | 0.93 |
| *Eugenia* sp. | 0 | 0 | 0 | 3.77 | 1.1 | 2.44 | 0 | 0 | 0 | 0 | 0 | 0 | 0.61 |
| *Malouetia tamaquarina* | 1.65 | 0.56 | 1.11 | 1.89 | 0.55 | 1.22 | 0 | 0 | 0 | 0 | 0 | 0 | 0.58 |
| *Protium* sp. | 1.65 | 0.37 | 1.01 | 1.89 | 0.55 | 1.22 | 0 | 0 | 0 | 0 | 0 | 0 | 0.56 |
| *Rinorea* sp.1 | 0.83 | 0.19 | 0.51 | 1.89 | 1.1 | 1.49 | 0 | 0 | 0 | 0 | 0 | 0 | 0.5 |
| *Aspidosperma carapanauba* | 0.83 | 0.19 | 0.51 | 1.89 | 0.55 | 1.22 | 0 | 0 | 0 | 0 | 0 | 0 | 0.43 |
| *Hymenopus heteromorphus* | 2.48 | 0.93 | 1.71 | 0 | 0 | 0 | 0 | 0 | 0 | 0 | 0 | 0 | 0.43 |
| *Jacaranda copaia* | 0.83 | 0.19 | 0.51 | 1.89 | 0.55 | 1.22 | 0 | 0 | 0 | 0 | 0 | 0 | 0.43 |
| *Patinoa paraensis* | 0.83 | 0.19 | 0.51 | 1.89 | 0.55 | 1.22 | 0 | 0 | 0 | 0 | 0 | 0 | 0.43 |
| *Pouteria* sp.1 | 2.48 | 0.56 | 1.52 | 0 | 0 | 0 | 0 | 0 | 0 | 0 | 0 | 0 | 0.38 |
| *Pachira aquatica* | 0 | 0 | 0 | 1.89 | 0.55 | 1.22 | 0 | 0 | 0 | 0 | 0 | 0 | 0.3 |
| *Pterocarpus officinalis* | 1.65 | 0.56 | 1.11 | 0 | 0 | 0 | 0 | 0 | 0 | 0 | 0 | 0 | 0.28 |
| *Maquira coriacea* | 1.65 | 0.37 | 1.01 | 0 | 0 | 0 | 0 | 0 | 0 | 0 | 0 | 0 | 0.25 |
| *Mouriri acutiflora* | 1.65 | 0.37 | 1.01 | 0 | 0 | 0 | 0 | 0 | 0 | 0 | 0 | 0 | 0.25 |
| *Ormosia coutinhoi* | 0.83 | 0.37 | 0.6 | 0 | 0 | 0 | 0 | 0 | 0 | 0 | 0 | 0 | 0.15 |
| *Ormosia* sp. | 0.83 | 0.37 | 0.6 | 0 | 0 | 0 | 0 | 0 | 0 | 0 | 0 | 0 | 0.15 |
| *Allantoma lineata* | 0.83 | 0.19 | 0.51 | 0 | 0 | 0 | 0 | 0 | 0 | 0 | 0 | 0 | 0.13 |
| *Cynometra cuneata* | 0.83 | 0.19 | 0.51 | 0 | 0 | 0 | 0 | 0 | 0 | 0 | 0 | 0 | 0.13 |
| *Guarea* sp. | 0.83 | 0.19 | 0.51 | 0 | 0 | 0 | 0 | 0 | 0 | 0 | 0 | 0 | 0.13 |
| *Guatteria* sp.2 | 0.83 | 0.19 | 0.51 | 0 | 0 | 0 | 0 | 0 | 0 | 0 | 0 | 0 | 0.13 |
| *Inga* sp. | 0.83 | 0.19 | 0.51 | 0 | 0 | 0 | 0 | 0 | 0 | 0 | 0 | 0 | 0.13 |
| *Platymiscium pinnatum* var. *ulei* | 0.83 | 0.19 | 0.51 | 0 | 0 | 0 | 0 | 0 | 0 | 0 | 0 | 0 | 0.13 |
| Sapotaceae sp. | 0.83 | 0.19 | 0.51 | 0 | 0 | 0 | 0 | 0 | 0 | 0 | 0 | 0 | 0.13 |
| *Spondias mombin* | 0.83 | 0.19 | 0.51 | 0 | 0 | 0 | 0 | 0 | 0 | 0 | 0 | 0 | 0.13 |
| *Tabebuia* sp.2 | 0.83 | 0.19 | 0.51 | 0 | 0 | 0 | 0 | 0 | 0 | 0 | 0 | 0 | 0.13 |
| *Vatairea guianensis* | 0.83 | 0.19 | 0.51 | 0 | 0 | 0 | 0 | 0 | 0 | 0 | 0 | 0 | 0.13 |
| **Total** | **100** | **100** | **100** | **100** | **100** | **100** | **100** | **100** | **100** | **100** | **100** | **100** | **100** |

**Fig. 4.**
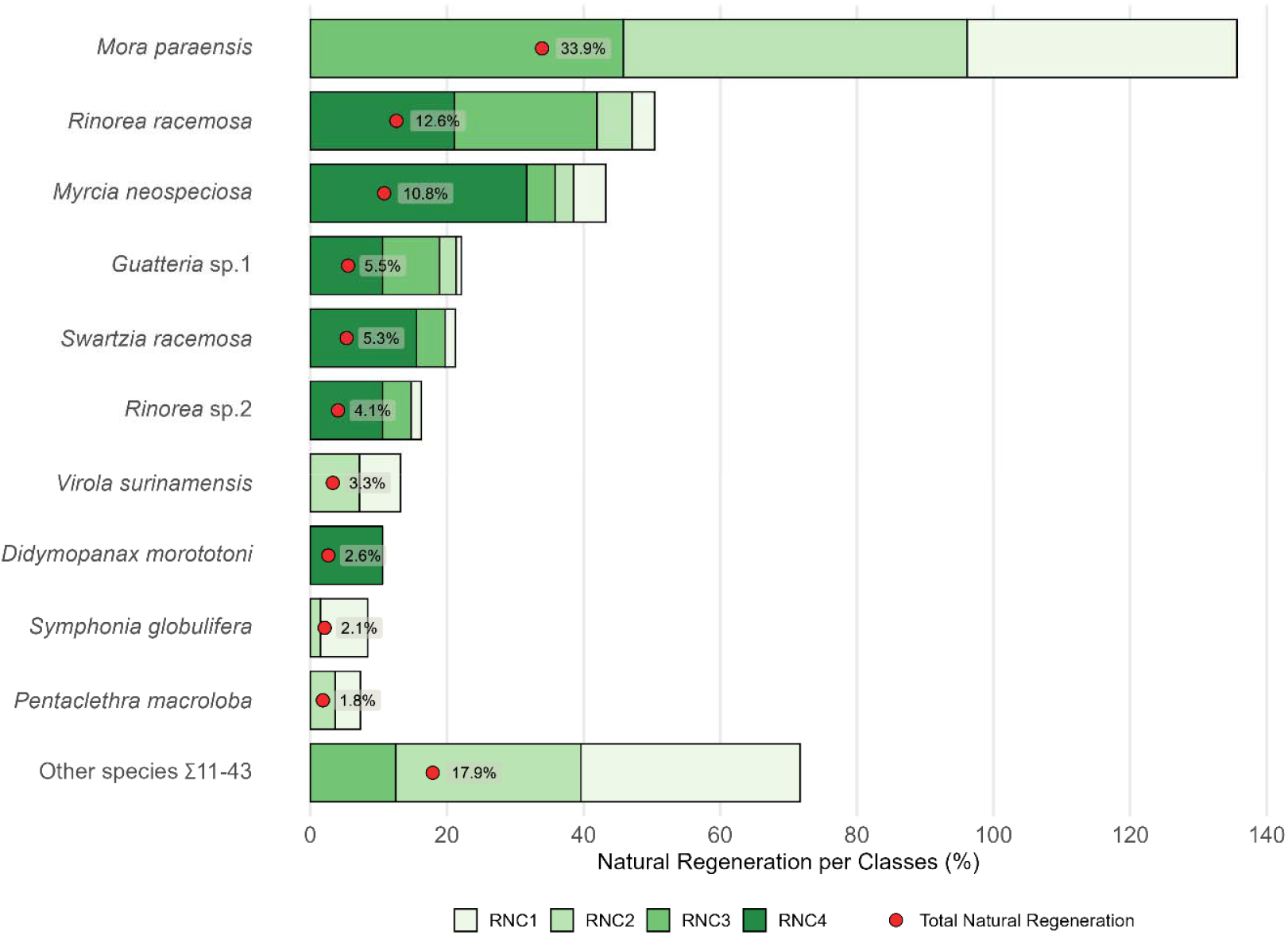
Ten species with the highest rates of Total Natural Regeneration (TNR) in decreasing order, including the values summed from remaining taxa in a tidal white-water floodplain forest of the Amazon River estuary, eastern Amazonia.

A total of 47% of species presented individuals in a single regeneration class (of which 85% were only first class), 35% of the total species occurred in two classes, 12% in three classes, and only 7% occurred in four regeneration classes evaluated. About 33% of the species showed individuals only in the first two height classes, such as *Virola surinamensis* and *Symphonia globulifera*. Despite being the most important species of this community, *Mora paraensis* was not represented in the fourth regeneration class. Only one species, *Didymopanax morototoni*, occurred in the fourth class. Most of the rare species occurred mainly in the first class.

The regeneration density recorded in the forest understory was 14,438 ind.ha^-1^, with species ranging from 19 to 9,695.2 ind.ha^-1^. When considering the inclusion criteria, the distribution frequency of sampled individuals was mainly concentrated in the first classes of DSH (70.5% of total individuals ≤ 1 cm) and height (70.8% of total individuals ≤ 100 cm) (Supplementary Information Fig. S3a, b). The total basal area amounted to 690.3 cm^2^ (or 1.3149 m^2^.ha^-1^), also presenting the highest number of individuals in the first classes (92.8% ≤ 2 cm^2^) (Supplementary Information Fig. S3c).

### Effect of distance from the river on regenerating tree species composition

The first two axes of the PCoA ordination captured a representative portion of the variation in species composition. For relative abundance data (i.e., proportional quantities of each species) and presence–absence data (i.e., only species occurrence, regardless of abundance), the explained variation was 39% and 44%, respectively. Species composition (PCoA-1) appeared to respond to the distance gradient from the Amazon River when relative abundance was considered (adj. r^2^ = 0.2179, F = 6.573, p = 0.019), showing a detectable shift along the gradient (Fig. 5a). In contrast, this relationship was not significant when based solely on presence–absence data (adj. r^2^ = 0.0478, F = 2.004, p = 0.173), indicating that variation in species occurrence was not strongly explained by river distance (Fig. 5b).

**Fig. 5.**
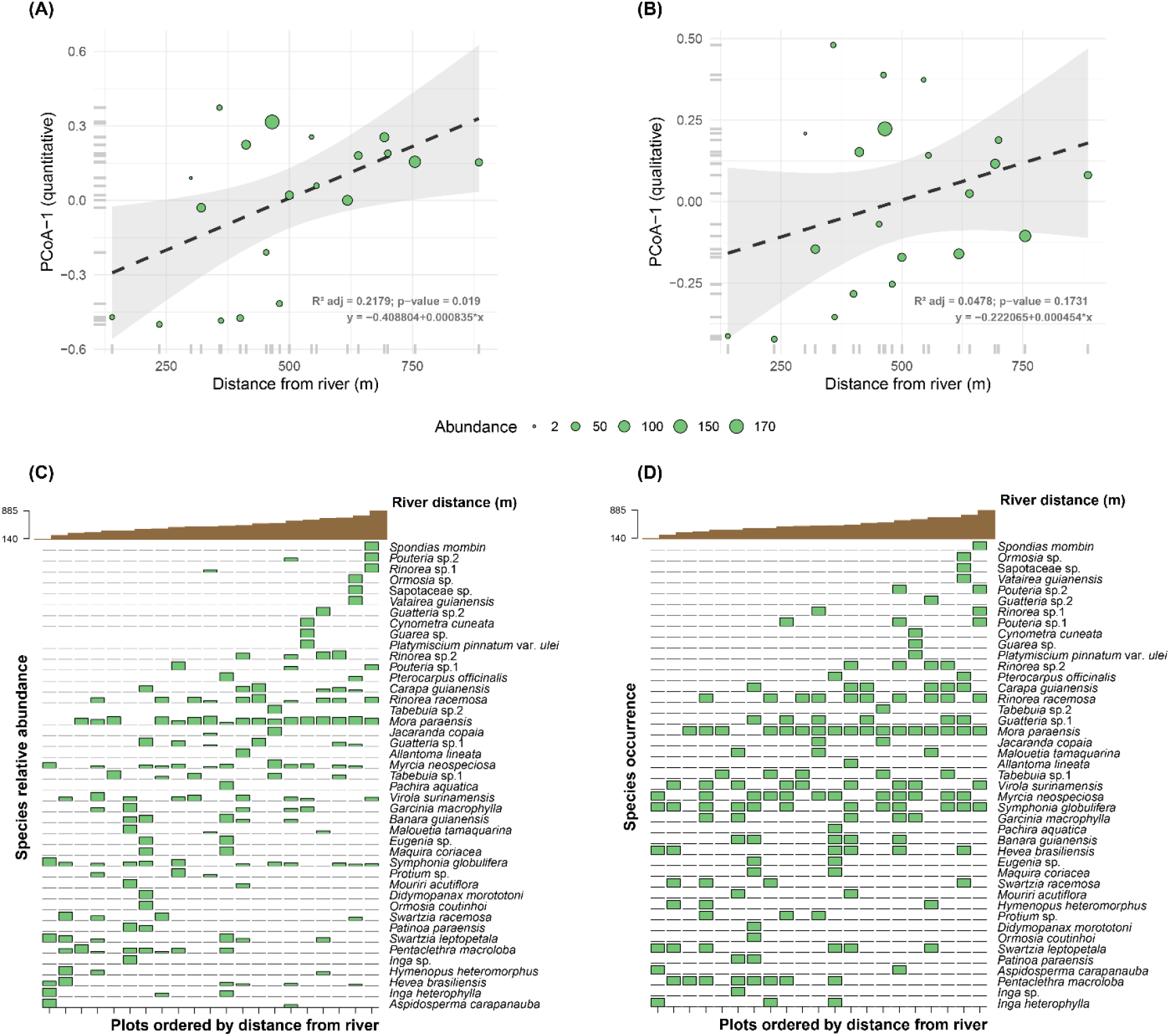
Relationship between species composition (PCoA-1) and river distance gradient for relative abundance (a) and presence-absence (b) data; each point represents a plot within the study area, and the proximity between them indicates the compositional similarity among plots. Variation in species composition along river distance gradient for relative abundance (c) and presence-absence (d) data, showing subtle changes in species composition from shorter to longer distances of the Amazon River mainstem.

The direct ordination based on relative abundance data shows that certain species dominate plots close to the main river channel but decline in importance as distance increases, while others become more abundant at sites farther from the river (Fig. 5c). A similar pattern is observed in the presence–absence data (Fig. 5d), indicating that most species occur across the entire gradient, although the abundance shifts detected (Fig. 5d) are not as clearly reflected in presence–absence. While some species appear more associated with areas near the river and others predominate in more distant areas, the five species identified as most abundant in the regenerating stratum (Fig. 3b) are also widely distributed along the distance gradient (Fig. 5d, d): *Mora paraensis, Virola surinamensis, Symphonia globulifera, Myrcia neospeciosa*, and *Rinorea guianensis*. In summary, regeneration is highly dominated by *M. paraensis* but is sustained by a suite of rare species that diversify the community.

## Discussion

We compiled a species list based on regenerating individuals in the APA Fazendinha, a small-scale survey conducted in an ecologically important tidal floodplain forest in the state of Amapá—a region that remains floristically underexplored compared to other parts of the Amazon Basin (Hopkins 2007; ter Steege et al. 2016; Stropp et al. 2020; Carvalho et al. 2023). This is the first study to document the composition (Table 1; Supplementary Information Fig. S2), diversity (Fig. 2), structure (Table 2; Figs. 3, 4; Supplementary Information Fig. S3), and distribution (Fig. 5) of regenerating assemblages in APA Fazendinha. These findings expand our understanding of species tolerant to tidal flood dynamics in Amazonian floodplain forests.

### Floristic diversity

The floristic profile of regenerating trees in the APA Fazendinha floodplain forest reflects a typical but strikingly oligarchic Amazonian pattern: very high species richness combined with extreme dominance. Fabaceae, represented by 12 species, emerged as the most important family, consistent with basin-wide patterns that identify legumes as both the most species-rich and ecologically versatile group across Amazonian forests and strata (ter Steege et al. 2006; Draper et al. 2021). Beyond their numerical dominance, Fabaceae are functionally central to floodplain systems (Wittmann et al. 2004, 2006; Assis et al. 2015; Feitosa et al. 2022), contributing traits such as symbiotic nitrogen fixation, ectomycorrhizal associations, and high seed mass, which promote both establishment under stressful edaphic conditions and competitive persistence (ter Steege et al. 2006, 2025; Feitosa et al. 2022).

Within this framework, *Mora paraensis* alone accounted for more than 70% of individuals and occurred in over 80% of plots (Table 2; Fig. 4), highlighting its role as a dominant species in the Amazon estuary. While its dominance is well-related for white-water floodplain forests (Wittmann et al. 2006; Albernaz et al. 2012; ter Steege et al. 2013; Carim et al. 2017), *M. paraensis* exemplifies a particularly narrow oligarchic strategy: unlike other dominants that are less restrictive geographically (e.g., *Eperua falcata, Mora excelsa*) (ter Steege et al. 2019), it is restricted to floodplains of the eastern Amazon and is less common in the central basin (Kubitzki and Ziburski 1994; Wittmann et al. 2013; Vasconcelos et al. unpubl. data). This geographic specialization suggests that its dominance is likely maintained more by local adaptations such as hydrocory-mediated dispersal (Cunha et al. 2017) and rapid germination during short exposure windows (Parolin 2009), than by broad ecological amplitude. The persistence of such a narrowly adapted dominant raises questions about its long-term resilience to environmental change, especially given its limited tolerance to prolonged submersion (Parolin 2012).

Other abundant species, including *Virola surinamensis, Symphonia globulifera, Myrcia neospeciosa*, and *Rinorea racemosa*, are recurrent elements of floodplain forests across Amazonia (Wittmann et al. 2006; Assis et al. 2015). Their widespread distribution across regeneration classes indicates greater demographic stability compared to the many rare taxa confined to initial stages. Together, these co-dominant species may provide functional redundancy, mitigating the ecological risks associated with *M. paraensis* monodominance. Overall, the consistent pattern of strong dominance of a small subset of tree species in neotropical forests is well-documented (MacArthur 1960; Brown 1984; Pitman et al. 2013; ter Steege et al. 2013, 2019; Williams et al. 2017), yet dominant trees display a variety of distributional patterns, even across different spatial scales. These species have been called “hyperdominant” at large geographical scales and “oligarchs” at regional-landscape scales when being abundant and frequent (Pitman et al. 2001; ter Steege 2013).

### Diversity estimates and sampling effort

The diversity metrics highlight a paradox typical of floodplain forests: low evenness but unexpectedly high Shannon diversity. Our Shannon index (H’ = 5.31) exceeds values commonly reported for floodplain forests in the estuarine region (2.5–3.6; Rabelo et al. 2000; Gama et al. 2002; Silva and Jardim 2016). This apparent inflation is explained by the coexistence of an oligarch species with a disproportionately large fraction of rare taxa (singletons and doubletons accounted for 53.5% of richness). Such a configuration elevates Shannon’s entropy because the index captures not only richness but also the presence of long “tails” of rare species.

Rarefaction and extrapolation curves further indicated that observed richness (43 species) underestimates the true community size, with at least nine additional species predicted under intensified sampling. This pattern echoes broader Amazonian findings where exhaustive sampling rarely exhausts species discovery due to high turnover at small spatial scales (Pitman et al. 2001; Stropp et al. 2020). Yet, even fourfold increases in sample size in similar environments have yielded diminishing returns in richness (Rabelo et al. 2000), suggesting that the floodplain community in APA Fazendinha appears to follow a saturation curve typical of oligarchic systems, where diversity stems less from local equitability and more from spatial heterogeneity and stochastic colonization events. Since determining the total number of species in an area is virtually impossible, especially in regions of exceptionally high richness, estimators are valuable tools for extrapolating observed richness and approximating total diversity from an incomplete community sample (Walther and Moore 2005).

### Regeneration structure

Regeneration density reached 14,438 ind.ha□^1^, heavily concentrated in the smallest size classes (≈70% of individuals ≤ 1 cm DSH or ≤ 100 cm height). The higher density of individuals in the smallest size classes is common in natural regeneration studies and represents a typical pattern of uneven-aged forests, as it usually results from the abundant production of propagules and recently established seedlings (Sccoti et al. 2011). This skewed structure is characteristic of uneven-aged tropical forests, where abundant recruitment is offset by high early mortality (Rother et al. 2013). Nearly half of the species were restricted to a single regeneration class (mostly the smallest), reflecting demographic bottlenecks that prevent many taxa from advancing to larger size categories. Such bottlenecks are likely driven by a combination of environmental filters (e.g., flood tolerance thresholds), microhabitat heterogeneity, and dispersal limitations.

In contrast, *M. paraensis, R. racemosa*, and *M. neospeciosa* exhibited high Total Natural Regeneration (TNR) across multiple classes, underscoring their capacity to shape the future canopy. These species exemplify a “structural insurance effect” (Silva et al. 2007), where continuous regeneration across stages may help ensure persistence even under disturbance. Conversely, *Carapa guianensis* and *Pentaclethra macroloba*, both of high socio-economic value in this region (Dantas et al. 2016, 2021), showed narrower class representation, which may compromise their long-term sustainability without management interventions. Recently, Dantas et al. (2022) documented a stable inverted “J” pattern for *P. macroloba* in the same protected area, indicating continuous regeneration into adult stages. This contrast suggests that local dynamics may differ from population-wide trends, and that even dominant and resilient species can experience recruitment constraints at finer scales. This underscores the importance of combining population-level and community-level perspectives to guide conservation and management.

Our findings further align with demographic analyses showing that *Mora paraensis* population growth appears to be particularly sensitive to survival of the smallest size classes rather than persistence of large adults (Fortini and Zarin 2011). This demographic profile helps explain why, despite intensive selective logging in the Amazon estuary, *M. paraensis* remains dominant in regeneration (this study, Miranda et al. 2018). If environmental conditions continue to favor seedling establishment, the species can maintain high recruitment rates and shows apparent demographic stability, even under pressure from timber extraction. The observed structure thus aligns with floodplain forests’ well-known tendency toward inequitable composition (Wittmann et al. 2004; Assis and Wittmann 2011), where a few species dominate both adult and regenerating strata, while many others persist only at low frequencies. This pattern highlights the dual ecological reality of tidal floodplains: simultaneously productive and fragile, resilient in some components but potentially vulnerable in others.

### River distance gradient

Although the relationship between river distance and regeneration composition was weak, the trend in abundance-based ordination suggests subtle structuring along the tidal gradient. This is consistent with the idea that presence-absence data may mask abundance shifts, especially in systems dominated by widely distributed oligarchic species. The weak gradient effect likely reflects the homogenizing influence of tidal hydrology, where twice-daily flood pulses redistribute propagules and impose recurrent submergence across plots (Junk et al. 2011; Cunha et al. 2017).

Oligarchic species tend to be structured by the same ecological mechanisms that create spatial heterogeneity in tree communities (Pitman et al. 2013). While they generally tolerate a wider range of environmental conditions than rarer taxa (Brown 1984), they still respond to local variation, often reaching peak abundances under the most favorable conditions (ter Steege et al. 2013). Thus, beta diversity may increase with environmental heterogeneity, but such variation is often driven more by shifts in oligarchic abundance than by changes in species identity.

Nevertheless, contrasting responses of co-occurring species, such as *P. macroloba* favoring lower sites and *C. guianensis* higher sites, point to the ecological importance of fine-scale topographic and edaphic variation (Pitman et al. 2001; Schietti et al. 2014; Tuomisto et al. 2017). These microhabitat filters, not captured by distance alone, likely structure regeneration niches in tidal floodplain forests. Future studies should incorporate direct measurements of elevation, soil texture, and flood duration to disentangle the relative roles of hydrology and edaphic heterogeneity.

### Methodological limitations and future directions

Despite the robust patterns identified, our study presents some methodological constraints that must be acknowledged. First, the identification of juvenile individuals remains a major challenge in Amazonian floristic surveys, particularly in hyperdiverse habitats where many taxa lack diagnostic traits at early life stages (Camargo et al. 2008). This may have led to conservative estimates of species richness. In addition, tropical forests comprise a few dominant and many rare tree species, but distinguishing the truly rare from those under-sampled remains a challenge for ecology and conservation. Second, we used distance from the mainstem river as a proxy for environmental gradients, but this variable only partially captures the fine-scale heterogeneity that likely drives regeneration dynamics, such as microtopography, soil fertility, and flood duration. Incorporating direct measurements of these variables would provide more accurate insights into the ecological filters shaping regeneration. Third, the rarefaction curve did not reach an asymptote, indicating that our sampling effort, although systematic, was insufficient to fully capture the species pool of the area. Increasing the number and spatial coverage of plots, and ideally integrating adult and regeneration strata, would allow stronger inferences about forest compositional continuity. Finally, our analyses reflect a static temporal snapshot of regeneration, whereas long-term monitoring would be necessary to capture demographic trajectories, successional shifts, and interannual variability associated with hydrological cycles.

### Conservation and management implications

Our results highlight both strengths and vulnerabilities of the APA Fazendinha floodplain forest. Despite intensive logging pressure (Fortini and Zarin 2011), the robust regeneration of *M. paraensis* ensures continuity of a key timber resource. On the other hand, the narrow dominance of this species raises concerns about resilience under climate change, as altered hydrological regimes could disproportionately impact its specialized life cycle. Furthermore, the persistence of many rare taxa only at early regeneration stages emphasizes the importance of conserving microhabitats that may serve as refugia for recruitment.

The dual structure of dominance and rarity highlights the importance of management strategies that protect both dominant stocks and the “long tail” of rare species. Monitoring regeneration dynamics, expanding floristic inventories (including adult strata), and integrating local ecological knowledge into conservation policies are essential steps for safeguarding estuarine floodplain forests. These ecosystems, already underrepresented in Amazonian research (Carvalho et al. 2023), may be especially vulnerable to synergistic pressures of deforestation, timber exploitation, and climate-driven changes (Flores et al. 2017; Stropp et al. 2020; Lapola et al. 2023).

## Conclusions

Our findings reveal a regenerating community dominated by a narrow oligarch, *Mora paraensis*, yet diversified by a high proportion of rare taxa. The simultaneous resilience through functional redundancy and vulnerability through demographic bottlenecks characterizes tidal white-water floodplain forests of the Amazon estuary. While *M. paraensis* recruitment suggests stability under current hydrological regimes, the fate of rare species underscores the importance of conserving microhabitat heterogeneity. Beyond providing a local baseline, this study raises broader hypotheses about how multiple flood pulses shape regeneration dynamics in Amazonian forests. We encourage long-term monitoring and multi-scale surveys to test these ideas, and to inform conservation strategies for estuarine floodplain forests, a neglected but ecologically pivotal ecosystem under growing anthropogenic and climatic pressures.

## Acknowledgements

The authors thank the Universidade do Estado do Amapá (UEAP) for providing the necessary infrastructure and the staff of Embrapa Amapá for providing some equipment used in field measurements. We also thank Nerivan Silva for assistance with species identification and valuable support during fieldwork.

## Author contributions

CCV, GF, CBS, CFC, JGLI, MFF, NLLA, VCCS, and JCA conceived the ideas and designed the methodology. CCV, GF, CBS, CFC, JGLI, MFF, NLLA, and VCCS conducted the fieldwork and identified the plant species. CCV analyzed the data with the support of GF, LLRP, KCMC, SCA, GGR, ARD, and RBL. CCV led the manuscript writing, later with significant input from GF, LLRP, and ARD. All authors contributed critically to the drafts and gave final approval for publication.

## Data availability

The original contributions presented in the study are included in the article/Supplementary Information. Further inquiries can be directed to the corresponding author.

## Declarations

### Ethical approval

Ethical approval was not applicable to this study.

