## Supplementary Information for "Tree species natural regeneration in a tidal white-water floodplain forest of the Amazon River estuary"

[1] Instituto Nacional de Pesquisas da Amazônia (INPA), Manaus, AM, Brazil
[2] Universidade do Estado do Amapá (UEAP), Macapá, AP, Brazil
[3] Universidade Federal do Espírito Santo (UFES), Jerônimo Monteiro, ES, Brazil
[4] Instituto de Pesquisas Científicas e Tecnológicas do Amapá (IEPA), Macapá, AP, Brazil
[5] Universidade Estadual do Maranhão (UEMA), São Luís, MA, Brazil
[6] Universidade Federal do Amapá (UNIFAP), Macapá, AP, Brazil
[7] Instituto Federal do Amapá (IFAP), Porto Grande, AP, Brazil
[8] Ação Ecológica Guaporé (ECOPORÉ), Porto Velho, RO, Brazil
[9] Instituto de Pesquisas Jardim Botânico do Rio de Janeiro (JBRJ), Rio de Janeiro, RJ, Brazil
[10] Embrapa Amapá, Macapá, AP, Brazil

*Corresponding author

**Table S1** List of R packages used in our analysis.

| Use | Package | Authorship | Identifier |
|---|---|---|---|
| **Data manipulation** | dplyr | Wickham et al. (2023) | https://doi.org/10.32614/CRAN.package.dplyr |
| | labdsv | Roberts (2023) | https://doi.org/10.32614/CRAN.package.labdsv |
| | readxl | Wickham et al. (2023) | https://doi.org/10.32614/CRAN.package.readxl |
| | tidyr | Wickham et al. (2024) | https://doi.org/10.32614/CRAN.package.tidyr |
| | writexl | Ooms et al. (2024) | https://doi.org/10.32614/CRAN.package.writexl |
| **Data analysis** | ape | Paradis et al. (2024) | https://doi.org/10.32614/CRAN.package.ape |
| | BiodiversityR | Kindt (2025) | https://doi.org/10.32614/CRAN.package.BiodiversityR |
| | iNEXT | Hsieh et al. (2024) | https://doi.org/10.32614/CRAN.package.iNEXT |
| | V.PhyloMaker | Jin and Qian (2019) | http://dx.doi.org/10.1111/ecog.04434 |
| | vegan | Oksanen et al. (2025) | https://doi.org/10.32614/CRAN.package.vegan |
| **Data visualization** | cowplot | Wilke (2024) | https://doi.org/10.32614/CRAN.package.cowplot |
| | ggplot2 | Wickham et al. (2024) | https://doi.org/10.32614/CRAN.package.ggplot2 |
| | ggpubr | Kassambara (2023) | https://doi.org/10.32614/CRAN.package.ggpubr |
| | ggtext | Wilke and Wiernik (2022) | https://doi.org/10.32614/CRAN.package.ggtext |
| | ggtree | Yu et al. (2017) | https://doi.org/doi:10.18129/B9.bioc.ggtree |
| | latex2exp | Meschiari (2022) | https://doi.org/10.32614/CRAN.package.latex2exp |

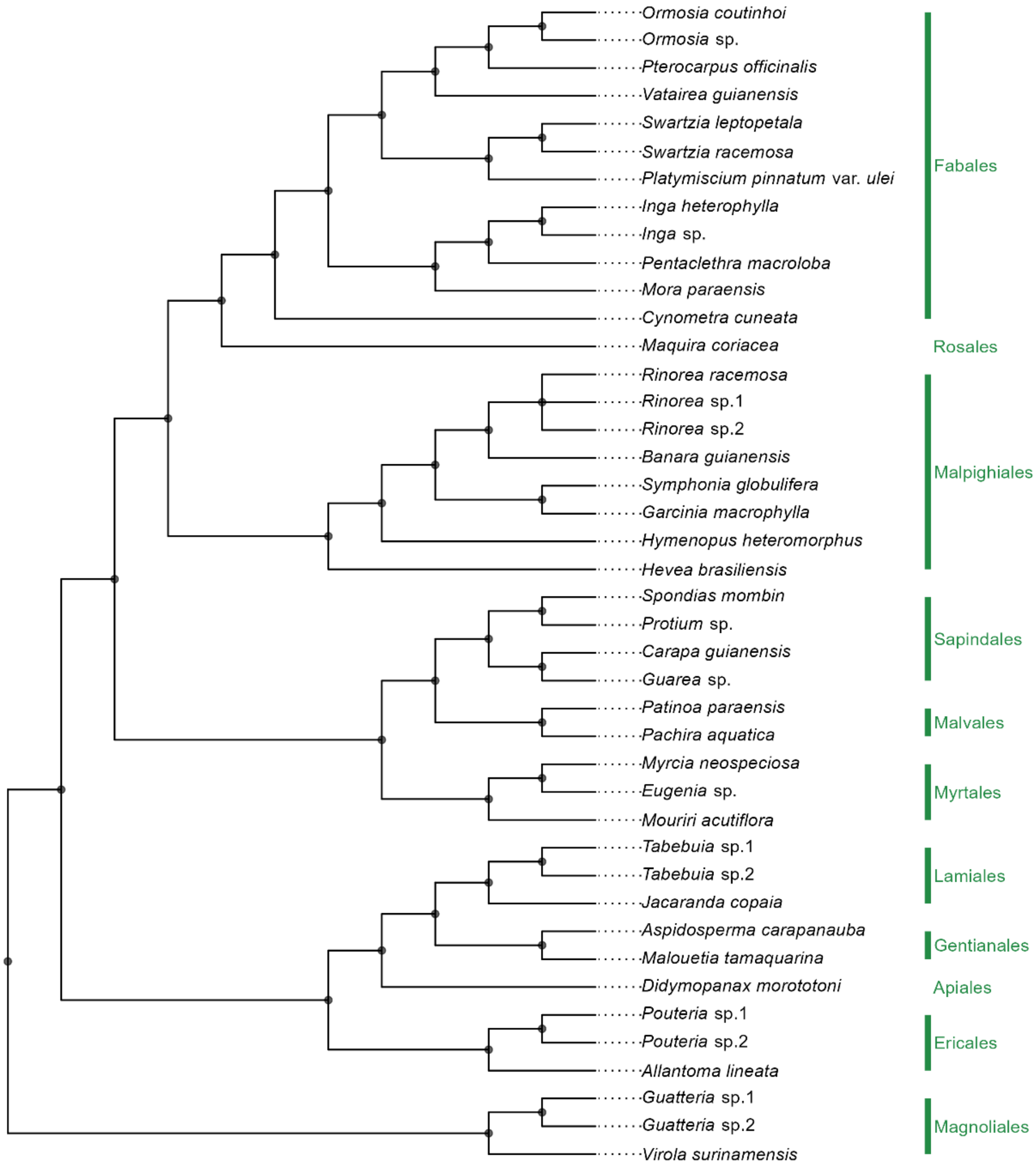


**Fig. S2** Representation of the phylogenetic relationships among the local species pool in a tidal white-water floodplain forest of the Amazon River estuary, eastern Amazonia. On the right side are the family and order to which each species belongs. The phylogenetic tree was generated with the 'S.PhyloMaker' function ('S' stands for seed plants) that generates phylogenies for seed plants based on the PhytoPhylo megaphylogeny (Qian and Jin 2016). Here, we used scenario three, which consists of adding genera or species to their families or genera using the same approach implemented in Phylomatic (Webb and Donoghue 2005) and BLADJ (A Simple Phylogenetic Branch Length Adjuster) (Webb 2000; Wikström et al. 2001).

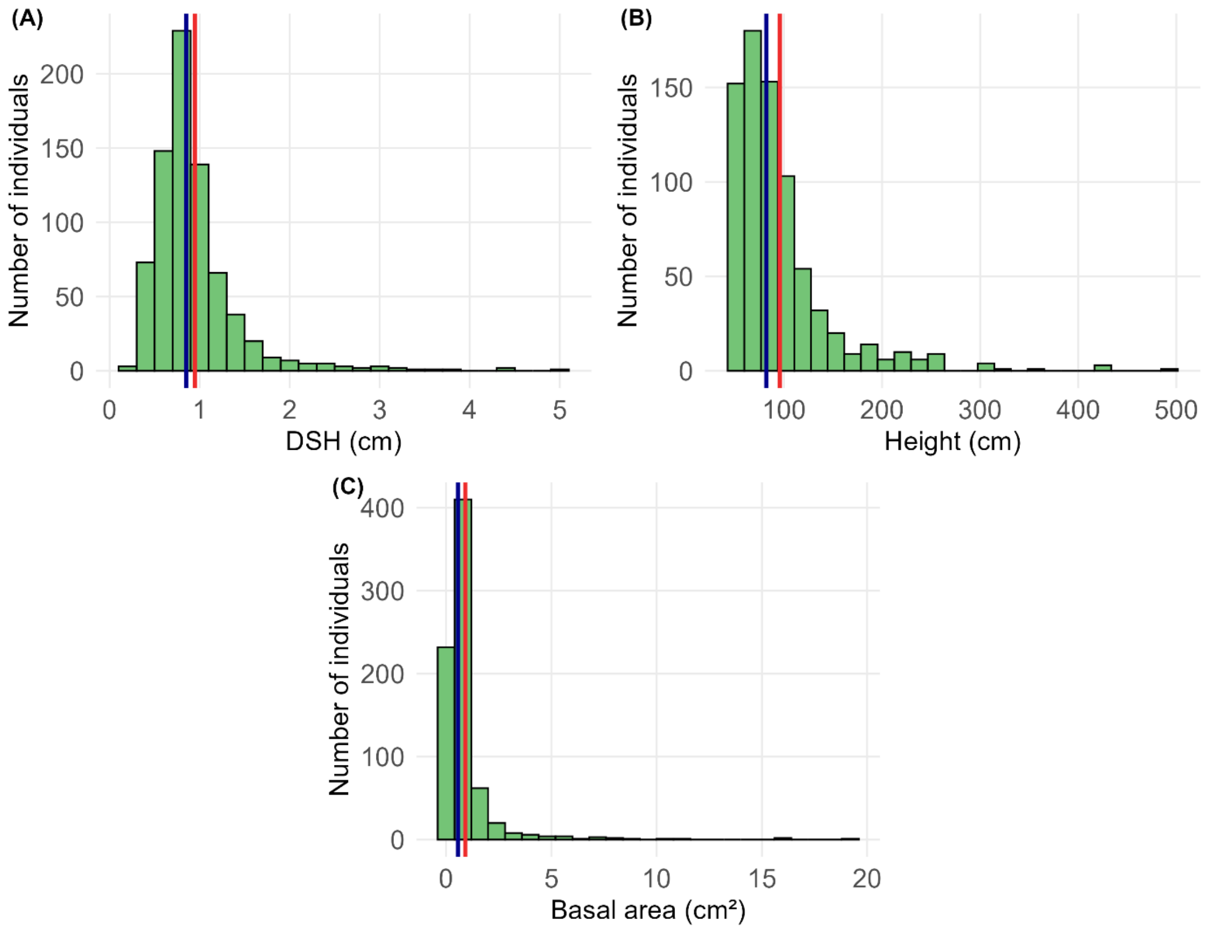


**Fig. S3** Frequency histograms showing the distribution of the number of individuals by DSH (Diameter at Soil Height) (a), height (b), and basal area ($cm^2$) (c) of the regenerating assemblage of tree species in a tidal white-water floodplain forest of the Amazon River estuary, eastern Amazonia. Vertical solid lines represent the position of the median (dark blue) and mean (red).
